# MRI-Visible Perivascular Space (PVS) Changes with Long-Duration Spaceflight

**DOI:** 10.1101/2021.08.26.457870

**Authors:** Kathleen E. Hupfeld, Sutton B. Richmond, Heather R. McGregor, Daniel L. Schwartz, Madison Luther, Nichole E. Beltran, Igor S. Kofman, Yiri E. De Dios, Roy F. Riascos, Scott J. Wood, Jacob J. Bloomberg, Ajitkumar P. Mulavara, Lisa Silbert, Jeffrey J. Iliff, Rachael D. Seidler, Juan Piantino

## Abstract

Humans are exposed to extreme environmental stressors during spaceflight and return with alterations in brain structure and shifts in intracranial fluids. To date, no studies have evaluated the effects of spaceflight on perivascular spaces (PVSs) within the brain, which are believed to facilitate fluid drainage and brain homeostasis. Here, we examined how the number and morphology of magnetic resonance imaging (MRI)-visible PVSs are affected by spaceflight, including prior spaceflight experience. Fifteen astronauts underwent six *T*_1_-weighted 3T MRI scans, twice prior to launch and four times following their return to Earth after ∼6-month missions to the International Space Station. White matter MRI-visible PVS number and morphology were calculated using an established automated segmentation algorithm. We found that novice astronauts showed an increase in total PVS volume from pre- to post-flight, whereas experienced crewmembers did not (adjusted for age, sex, and time between landing and first MRI scan). Moreover, experienced astronauts exhibited a significant correlation between more previous flight days and greater PVS median length at baseline, suggesting that experienced astronauts exhibit holdover effects from prior spaceflight(s). There was also a significant positive correlation between pre- to post-flight increases in PVS median length and increases in right lateral ventricular volume. The presence of spaceflight associated neuro-ocular syndrome (SANS) was not associated with PVS number or morphology. Together, these findings demonstrate that spaceflight is associated with PVS morphological changes, and specifically that spaceflight experience is an important factor in determining PVS characteristics.

## 1. Introduction

Spaceflight-induced physiological stressors have been well documented, including exposure to microgravity, ionizing radiation, and circadian disruption^1,2^. Astronauts returning to Earth after spaceflight exhibit structural brain changes, including an upward shift of the brain within the skull^3–5^, regional gray matter volume changes^4–7^, and altered white matter volume and microstructural integrity^6–8^. Spaceflight is also associated with alterations in cerebrospinal fluid (CSF) flow, evidenced by ventricular expansion^3,5–7,9–12^ and redistribution of CSF away from the top of the brain^3,6,8^. Clinically, approximately 33% of astronauts returning from long-duration spaceflight exhibit spaceflight associated neuro-ocular syndrome (SANS), a constellation of ocular structural changes including optic disc edema, choroidal folding, globe flattening, and hyperopic refractive error shifts^13,14^. The etiology of SANS remains unknown but could include fluid shifts, increased intracranial pressure, and / or altered CSF drainage with spaceflight^13^.

Brain CSF^3,5,8,9,12^, gray^4,5^, and white matter^8,12^ changes with spaceflight are associated with the number and duration of prior missions. Furthermore, some brain changes (particularly, ventricular expansion) show only partial recovery at 6 to 12 months post-flight^5,6,10,11^; thus, astronauts who return to space may experience holdover effects from prior flights. For instance, Hupfeld and colleagues^5^ reported that a shorter time between missions was associated with larger pre-flight ventricles (even after correcting for age effects). Moreover, they found that experienced astronauts who exhibited *larger* pre-flight ventricles also showed *less* ventricular expansion in a subsequent flight compared to novice crewmembers^5^. Together, these findings highlight both frequency and duration of spaceflight as important factors for understanding how microgravity affects cranial fluid shifts and CSF drainage.

According to the classic model, CSF produced by the choroid plexus circulates to the subarachnoid space before exiting the brain via the arachnoid granulations. However, recent evidence suggests that CSF in the subarachnoid space also flows into the brain parenchyma through the periarterial spaces surrounding the penetrating arteries^15^. Once in the brain parenchyma, CSF exchanges with interstitial space fluid (ISF), eventually returning to the cisternal CSF along the perivenous pathway^16^. Interstitial and CSF solutes are cleared from the cranium in part through the cervical lymphatic system via dural lympahtic vessels located along the superior sagittal sinus, or near the cribriform plate^17^. This perivascular exchange of CSF and ISF along what has been termed the glymphatic pathway plays an important role in the removal of cerebral interstitial solutes and wastes^18,19^. Perivascular space (PVS) enlargement visible on MRI is proposed to represent a disruption of this process, and has been observed in conditions associated with glymphatic dysfunction^20–25^. To date, the effect of microgravity on perivascular CSF circulation remains unknown. We hypothesized that the upward cerebral shift resulting from microgravity exposure compresses the meningeal lymphatics running along the superior sagittal sinus, impairing CSF/ISF exchange. Disruption of fluid drainage would be evidenced by an increase in PVS number and / or changes in the morphological aspects of MRI-visible PVS (i.e., their volume, length, or width).

The objectives of this study included: 1) To define the effects of spaceflight on PVS characteristics. We predicted an increase in white matter PVS number, volume, width, and / or and / or length after flight. 2) To explore the differences in PVS changes with flight for novice versus experienced astronauts. We predicted that novice astronauts would exhibit more changes in PVS number and size from pre- to post-flight. We also predicted that, within the experienced astronaut subgroup, there would be a correlation between a greater number of days previously spent in spaceflight and a higher number and larger size of PVSs at baseline. 3) To determine the relationship between ventricular expansion and PVS changes with spaceflight. We predicted that greater pre- to post-flight increases in ventricular volume would be associated with an increase in PVS number and size. 4) To examine differences in PVS changes with flight for those who exhibited signs of SANS vs. those who did not (no-SANS). We predicted greater increases in PVS number and size from pre- to post-flight for the SANS versus no-SANS individuals.

## 2. Methods

### 2.1 Participants

This was a longitudinal cohort study. Fifteen astronauts provided their written informed consent to participate in this study. The University of Michigan, University of Florida, and NASA Institutional Review Boards approved all study procedures. This analysis was implemented as part of a larger NASA-funded project investigating brain^5,26^ and behavioral^27^ changes occurring with long-duration ISS missions^28^.

### 2.2 Experimental Design

The astronauts completed MRI scans at six time points: approximately 180 and 60 days prior to launch (i.e., Launch-180 and Launch-60 days), as well as approximately 4, 30, 90, and 180 days after return to Earth (i.e., Return+4, Return+30, Return+90, and Return+180 days, respectively). One astronaut withdrew from the study before their Return+180 days session; thus, we acquired data from 14 of 15 astronauts at this time point.

### 2.3 Structural MRI Acquisition

All MRI scans were collected using the same 3.0 Tesla Siemens Magnetom Verio MRI scanner at the University of Texas Medical Branch at Victory Lakes, TX. We collected a *T*_1_- weighted structural MRI scan using the following imaging parameters: magnetization-prepared rapid gradient-echo (MPRAGE) sequence, TR = 1.9 s, TE = 2.32 ms, flip angle = 9°, FOV = 250 × 250 mm, slice thickness = 0.9 mm, 176 sagittal slices, matrix = 512 × 512, voxel size = 0.488 × 0.488 × 0.9 mm = 0.214 mm^3^.

### 2.4 Measurement of Right Lateral Ventricular Volume

We first processed all *T*_1_-weighted scans using the Computational Anatomy Toolbox (CAT12.6, version 1450)^29,30^ implemented within Statistical Parametric Mapping 12 (SPM12; version 7219)^31^ using MATLAB (R2016a, MathWorks Inc., Natick, MA). We implemented standard CAT12 preprocessing steps to extract total white matter and total intracranial volume, as well as native space right lateral ventricular volume for each participant and time point, defined by the Neuromorphometrics volume-based atlas map (http://Neuromorphometrics.com).

We used right ventricular volume in all statistical models because previous work by our group demonstrated a right-biased lateral ventricular enlargement in head-down-tilt bed rest participants who experienced more severe ocular pathology (i.e., SANS symptoms)^32^. Further justification for focusing on right-side lateral ventricular volume includes that CSF predominantly drains on the right side^33,34^, and optic disc edema in astronauts occurs more prominently on the right side^35^.

### 2.5 Characterization of PVSs

#### 2.5.1 PVS Segmentation

White matter PVS identification was carried out using our previously-described algorithm that searches for hypointense structures that meet certain pre-specified morphological criteria^36–38^. Briefly, first, the cerebellum, basal ganglia, and midbrain of the 3D *T*_1_-weighted MRI were masked (via FreeSurfer (Version 5.1)), and the images were skull-stripped using the Brain Extraction Tool (BET) from the FMRIB Software Library (FSL; Version 5.0). Resultant images were then segmented via FreeSurfer into four tissue types: white matter, cortical gray matter, subcortical gray matter, and ventricular CSF. Next, we corrected white matter masks for any tissue misclassification and eroded by one voxel to avoid partial volume effects, as described by Promjunyakul et al.^39^ The PVSs automatically detected by our algorithm were manually verified by a single, trained rater (ML). Any identified false alarms that were not PVS were removed from the final statistical analyses, as done in our prior work^36,37^. We then used custom MATLAB (R2016b; MathWorks Inc., Natick, MA) code to calculate five key outcome metrics, as done in our past work^36–38^. These outcomes included two metrics to describe total PVS: 1) total PVS volume (i.e., the sum of the volumes in mL of each distinct PVS in the white matter), and 2) total PVS number (i.e., the total number of distinct PVS in the white matter). These outcome metrics also included three which described characteristics of each distinct PVS: volume (mL), length (mm), and width (mm). For these three metrics, we then calculated the median volume, median length, and median width across all PVS clusters for each participant at each time point.

#### 2.5.2 Image Quality Control

Out of the 89 total scans (i.e., 6 time points × 15 astronauts, with one astronaut who withdrew before the final post-flight scan), we excluded four scans in total from further analysis of PVS metrics as these scans were degraded by motion artifact. These included only one pre-flight scan (i.e., Launch-60 days); for this participant, we calculated the percent change in PVS metrics based on their only baseline scan (i.e., Launch-180 days). No scans from the first post-flight time point were excluded. The other three excluded scans were from later post-flight timepoints (i.e., one from Return+90 days, and two from Return+180 days).

### 2.6 Statistical Analyses

We conducted all statistical analyses using R 4.0.0^40^.

#### 2.6.1 Baseline Reliability of PVS Metrics

We first tested for a stable baseline in the five PVS metrics and right lateral ventricular volume by calculating the intraclass correlation coefficient (ICC) between the two pre-flight scans^41^. As in our previous work^42^, we planned *a priori* to exclude from further statistical analyses any metric with an ICC(3,*k*) value below 0.5 (moderate reliability^43^).

#### 2.6.2 PVS Changes with Spaceflight

Next, we used five separate multivariate linear models to test for changes with spaceflight in each PVS metric and right lateral ventricular volume. We entered the percent change in the PVS metric or ventricular volume from the last pre-flight time point (Launch-60 days) to the first post-flight time point (Return+4 days) as the outcome variable, and included the following covariates: sex, mean-centered age at launch, and mean-centered time elapsed from landing to the Return+4 days MRI scan. In this case, we were interested in whether the intercept was significant (*p* < 0.05), thereby indicating a significant change in the PVS metric or ventricular volume with spaceflight. We then reran these models also including novice versus experienced status as a covariate, to test for differences in pre- to post-flight PVS and ventricular volume changes based on prior flight exposure. We ensured that each model met the relevant assumptions (i.e., independence, outliers, linearity, and normality). We tested for normality using visual inspection of qqplots and the Shapiro test (*p* > 0.05).

#### 2.6.3 Relationship Between Pre-Flight PVS Characteristics and Previous Flight Experience

Next, we conducted an exploratory analysis on the subset of *n* = 6 experienced astronauts. We used stepwise multivariate linear regression to examine whether average baseline PVS metrics or right lateral ventricular volume associated with the total number of previous days spent in space (i.e., total days in space, not including the current mission). We first ran a reduced model including previous flight days as the only predictor. If there was a significant (*p* < 0.05) linear relationship between previous flight days and the respective PVS metric, we calculated a full model including sex and mean-centered age at launch as additional covariates. We then used stepAIC^44^ to produce a final model that retained only the best predictor variables; stepAIC selects a maximal model based on the combination of predictors that produces the smallest Akaike information criterion (AIC). As above, we ensured that each model met the relevant assumptions, and we tested for normality using visual inspection of qqplots and the Shapiro test (*p* > 0.05). As these models examined baseline differences (instead of percent change with flight), to account for individual differences in total brain tissue volumes and allow for direct inter-subject comparison of PVS metrics, in these models we expressed each participant’s total PVS volume and number at baseline as a percentage of their average baseline total brain white matter volume. To account for individual differences in head size and allow for direct inter-subject comparison of ventricular volume, in these models we expressed each astronaut’s right lateral ventricular volume at baseline as a percentage of their average baseline total intracranial volume. A correction for head size was not necessary for the median PVS metrics.

#### 2.6.4 Relationship Between Spaceflight Changes in PVS Characteristics and Ventricular Volume

We also used stepwise multivariate linear regression to examine whether pre- to post-flight percent change in right lateral ventricular volume predicted changes in PVS characteristics. As there were no differences in ventricular volume changes with flight for the novice versus experienced astronauts, we conducted this analysis across the entire (*n* = 15) astronaut cohort. We first ran a reduced model including right lateral ventricular volume as the only predictor. If there was a significant (*p* < 0.05) linear relationship between ventricular volume and the respective PVS metric, we calculated a full model including sex and mean-centered age at launch, time elapsed from landing to the Return+4 days MRI scan, and total flight duration as additional covariates. Then, as above, we used stepAIC^44^ to produce a final model that retained only the best predictor variables, based on AIC. As above, we ensured that each model met the relevant assumptions (i.e., independence, outliers, linearity, and normality), and we tested for normality using visual inspection of qqplots and the Shapiro test (*p* > 0.05).

#### 2.6.5 Differences in PVS Changes for SANS versus no-SANS Subgroups

We conducted a final exploratory analysis comparing a subset of astronauts who developed at least one symptom of spaceflight associated neuro-ocular syndrome (SANS, *n* = 6) to the subset of astronauts who did not experience any SANS symptoms (no-SANS, *n* = 6). SANS status was determined using pre- and post-flight ophthalmic exam results obtained from the NASA Lifetime Surveillance of Astronaut Health Repository. We were unable to obtain SANS data for three astronauts, resulting in a sample of 12 for our SANS analysis. As recently defined by Lee et al. 2020^13^, an astronaut was determined to have SANS if they showed any of the following change in either eye from pre- to post-flight: optic disc edema (variable Frisén grades), choroidal folds, hyperopic refractive error shifts > 0.75 D, or globe flattening. We conducted identical analyses to those described above (Section 2.6.2) to test for pre- to post-flight change in the PVS metrics and right lateral ventricular volume, but here we included SANS status as a predictor of interest instead of novice versus experienced status.

## 3. Results

The cohort consisted of 15 astronauts, 9 of whom had no prior flight experience (“novice” astronauts) and 6 of whom had completed at least one previous spaceflight mission (“experienced” astronauts). The experienced astronauts were, on average, older than the novice astronauts, thus we controlled for age at launch in all statistical models. No other group differences emerged for sex, mission duration, or time between landing and the first post-flight MRI scan (Table 1).

**Table 1.** Astronaut Demographics and Flight Experience

Note.* \* $p < 0.05$ , \*\* $p < 0.01$ .
| | Novice | Experienced | t or $\chi^2$ | p |
| --- | --- | --- | --- | --- |
| **Demographics** |  |  |  |  |
| Number of astronauts | 9 | 6 | -- | -- |
| Age at Launch (years) | 43.98 (4.90) | 52.67 (4.18) | 3.68 | **0.003\*\*** |
| Sex | 33% F | 17% F | 0.51 | 0.475 |
| **Current Mission** |  |  |  |  |
| Mission duration (days) | 166.78 (31.37) | 226.50 (71.21) | 1.93 | 0.099 |
| Time elapsed between landing and Return+4 days MRI scan | 4.83 (0.41) | 4.33 (1.41) | 1.00 | 0.341 |
| **Experienced Astronauts (n = 6)<sup>a</sup>** |  |  |  |  |
| Number of previous missions | -- | 2.00 (0.89) | -- | -- |
| Past flight days (days)<sup>b</sup> | -- | 187.50 (151.67) | -- | -- |
| Time elapsed since last flight (years)<sup>c</sup> | -- | 5.77 (1.60) | -- | -- |

<sup>a</sup> Here, we report further details regarding previous spaceflight experience for $n = 6$ of 15 astronauts who completed at least one flight before the current ISS mission.
<sup>b</sup> Past flight days indicates the total number of previous days spent in flight, not including the current mission.
<sup>c</sup> Time elapsed since the last flight is the time from the landing day of the most recent flight to the launch of the current mission.

### 3.1 Identification of PVSs

We identified PVSs in each astronaut at each time point. PVSs were identified in the white matter in various regions across the whole brain (see exemplar participant, Figure 1).

**Fig 1.**
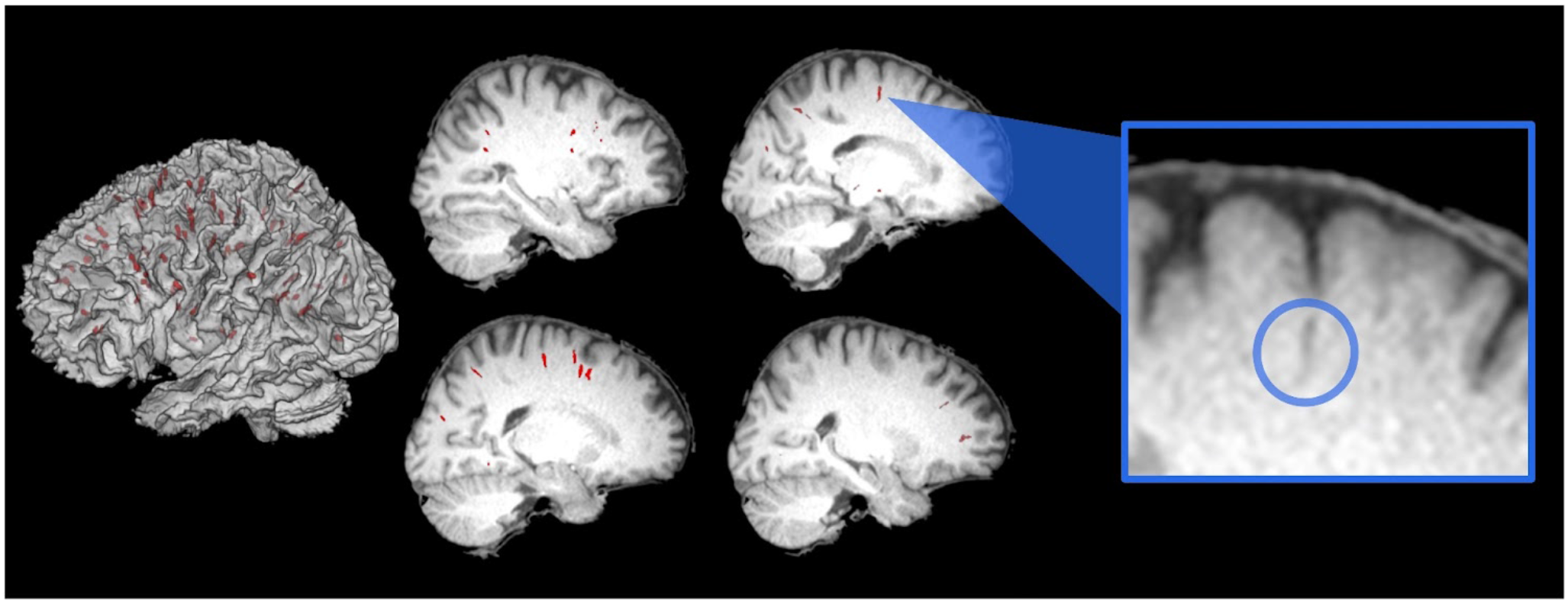
PVSs Identified on a Single Astronaut. Here we depict a binary mask of PVSs for a single astronaut at the last pre-flight time point (Launch-60 days), for illustrative purposes. PVSs are shown in red, overlaid onto this individual’s native space 3D-rendered white matter segment (*left*) as well as several sagittal slices of their skull stripped native space structural scan (*right*). The blue box shows a zoomed-in view of one PVS, inside the blue circle.

### 3.2 Baseline Reliability of PVS Metrics

Four of the five PVS metrics, as well as right lateral ventricular volume, yielded baseline ICC(3,*k*) values > 0.50. ICC (3,*k*) values were as follows: <u>total PVS volume</u>: 0.98, <u>total PVS number</u>: 0.97, <u>median PVS volume</u>: 0.58, <u>median PVS length</u>: 0.52, <u>median PVS width</u>: 0.38, and <u>right lateral ventricular volume</u>: 1.00. We excluded median width from further statistical analyses due to the low pre-flight reliability of this measure.

### 3.3 Changes in PVS Characteristics with Spaceflight

Changes in PVS characteristics with spaceflight are summarized in Table S1, Table 2, and Fig. 2. There were no significant whole-group changes in any of the PVS characteristics from pre- to post-flight (*p* > 0.05; Table S1). However, compared to their experienced colleagues, after adjusting for age, sex, and time between landing to the first MRI, the novice astronauts showed a mean pre- to post-flight increase in total PVS volume (*p* = 0.028; Table 2 shows the model outputs including covariate effects, while Fig. 2 shows unadjusted percent changes by the novice versus experienced subgroups). There was no pre-to post-flight difference by novice versus experienced status for the other PVS characteristics.

**Table 2.**
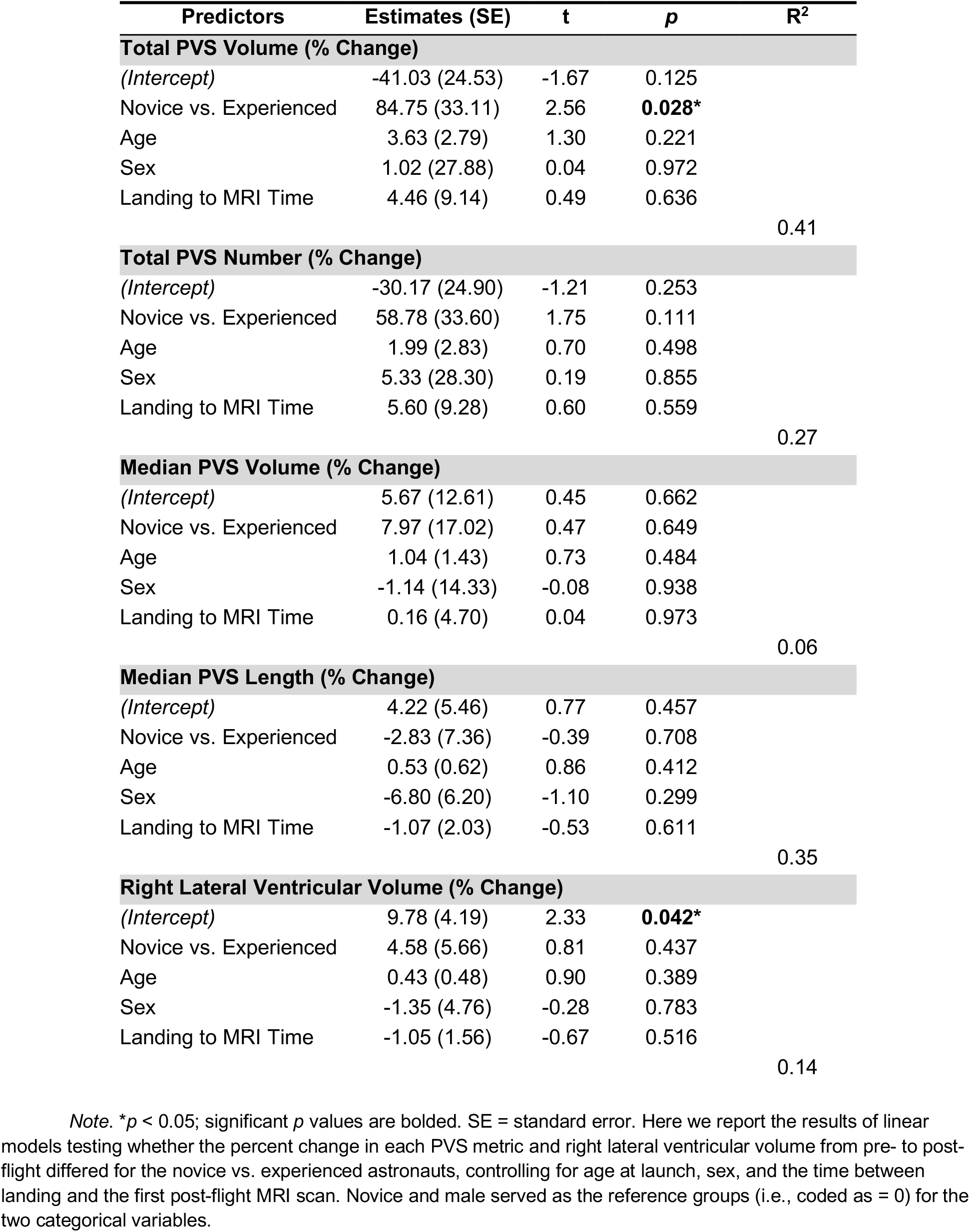
PVS Changes and Ventricular Expansion from Pre- to Post-Flight: Novice vs. Experienced Differences

**Fig 2.**
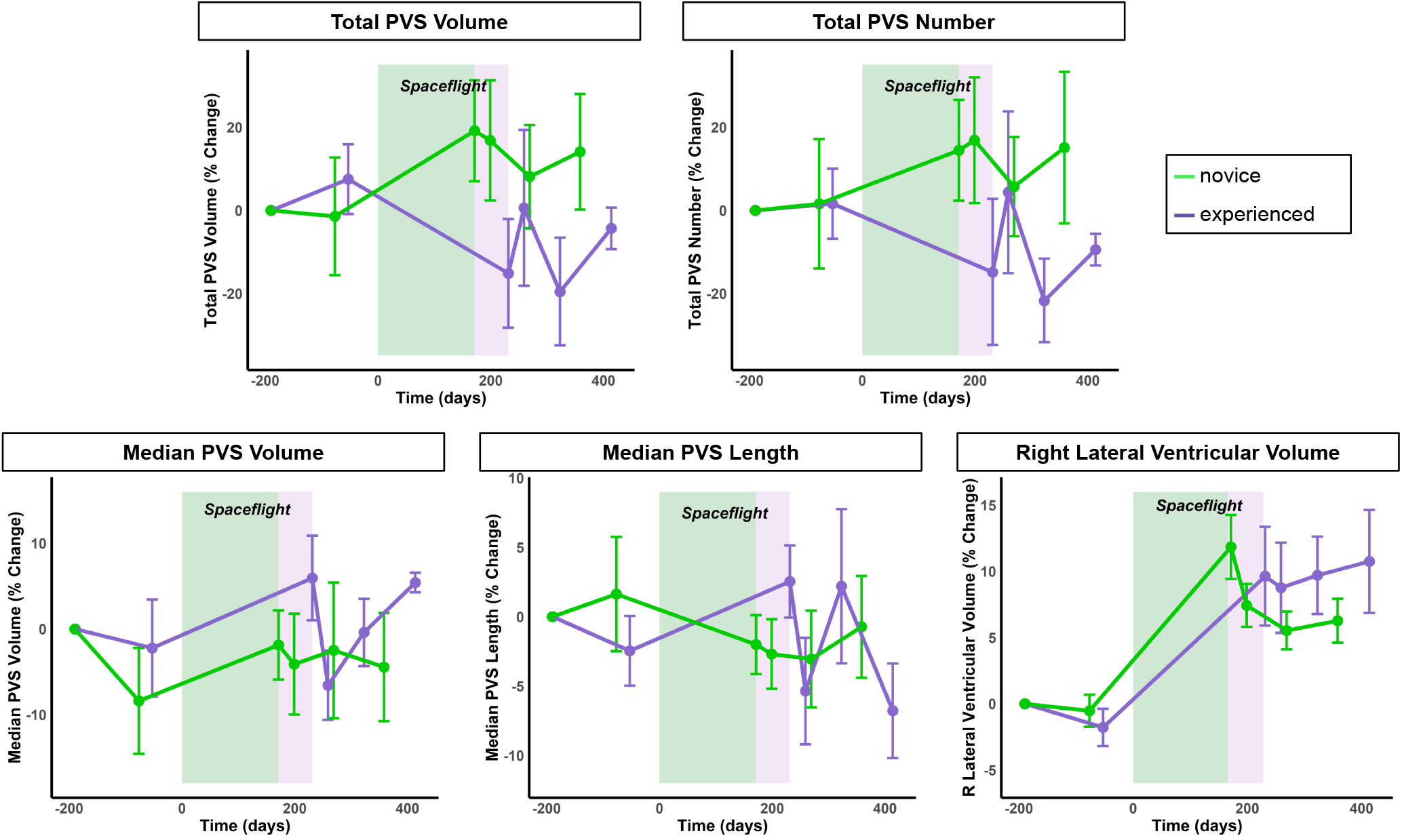
Changes in PVS Metrics and Right Lateral Ventricular Volume from Pre- to Post-Flight. Group average PVS characteristics (all expressed as % change from the first scan) are depicted for each of the six time points, and split by novice (green) and experienced (purple) astronauts. Bars represent standard error. The width of the green and purple boxes indicates the average flight duration for novice and experienced astronauts, respectively.

After adjusting for the above-mentioned covariates, right ventricular volume showed a significant increase from pre- to post-flight for the whole group (*p* = 0.0002; Table S1), though there was no effect of experience level on ventricular size changes (*p* > 0.05; Table 2).

### 3.4 Correlation Between Past Flight Experience and Baseline PVS Characteristics

The correlations between past flight experience and baseline PVS characteristics are summarized in Table 3 and Fig. 3 (Table 3 shows all model effects, whereas unadjusted values are plotted in Fig. 3 for ease of interpretation). Among the experienced astronauts (*n* = 6), number of previous flight days was correlated with pre-flight PVS median length. The stepwise variable selection procedure retained both previous days in space (*p* = 0.003) and age at launch (*p* = 0.025) as significant predictors of baseline PVS median length. Controlling for age, for every day previously spent in space (not including the current mission), there was an average 0.01 mm increase in PVS length (*p* = 0.003). There was no correlation between number of previous flight days and the remaining pre-flight PVS characteristics or ventricular volume.

**Table 3.** Relationship of Pre-Flight PVS Characteristics and Ventricular Volume with Past Flight Experience

Note.* \* $p < 0.05$ , \*\* $p < 0.01$ , \*\*\* $p < 0.001$ ; significant $p$ values are bolded.
| Predictors | Estimates (SE) | t | p | R<sup>2</sup> |
| --- | --- | --- | --- | --- |
| **Total PVS Volume (Average Baseline, % of Total Brain White Matter Volume)<sup>a</sup>** |  |  |  |  |
| (Intercept) | 0.13 (0.12) | 1.07 | 0.362 |  |
| Past Flight Days | 0.002 (0.001) | 1.67 | 0.193 |  |
| Age | -0.05 (0.04) | -1.08 | 0.358 |  |
|  |  |  |  | 0.55 |
| **Total PVS Number (Average Baseline, % of Total Brain White Matter Volume)<sup>a</sup>** |  |  |  |  |
| (Intercept) | 3.44 (2.04) | 1.69 | 0.190 |  |
| Past Flight Days | 0.04 (0.02) | 1.93 | 0.149 |  |
| Age | -0.95 (0.70) | -1.35 | 0.270 |  |
|  |  |  |  | 0.60 |
| **Median PVS Volume (Average Baseline, mm<sup>3</sup>)** |  |  |  |  |
| (Intercept) | 31.64 (4.00) | 7.92 | **0.004\*\*** |  |
| Past Flight Days | 0.05 (0.04) | 1.38 | 0.263 |  |
| Age | -0.91 (1.38) | -0.66 | 0.555 |  |
|  |  |  |  | 0.54 |
| **Median PVS Length (Average Baseline, mm)** |  |  |  |  |
| (Intercept) | 10.54 (0.11) | 96.98 | **< 0.001\*\*\*** |  |
| Past Flight Days | 0.01 (0.001) | 8.81 | **0.003\*\*** |  |
| Age | -0.16 (0.04) | -4.17 | **0.025\*** |  |
|  |  |  |  | 0.98 |
| **Right Lateral Ventricular Volume (Average Baseline, % of Total Intracranial Volume)<sup>b</sup>** |  |  |  |  |
| (Intercept) | 0.53 (0.14) | 3.69 | **0.034 \*** |  |
| Past Flight Days | 0.001 (0.001) | 1.02 | 0.383 |  |
| Age | -0.06 (0.05) | -1.20 | 0.316 |  |
|  |  |  |  | 0.33 |

<sup>a</sup> To account for individual differences in total brain tissue volumes, total PVS volume and number at baseline were normalized as follows: (total PVS volume (mL) or number) / (average pre-flight total brain white matter volume (mL))\*100%.
<sup>b</sup> To account for individual differences in total brain volume, right lateral ventricular volume at baseline was normalized as follows: (right lateral ventricular volume (mL)) / (average pre-flight total intracranial volume (mL))\*100%.

**Fig 3.**
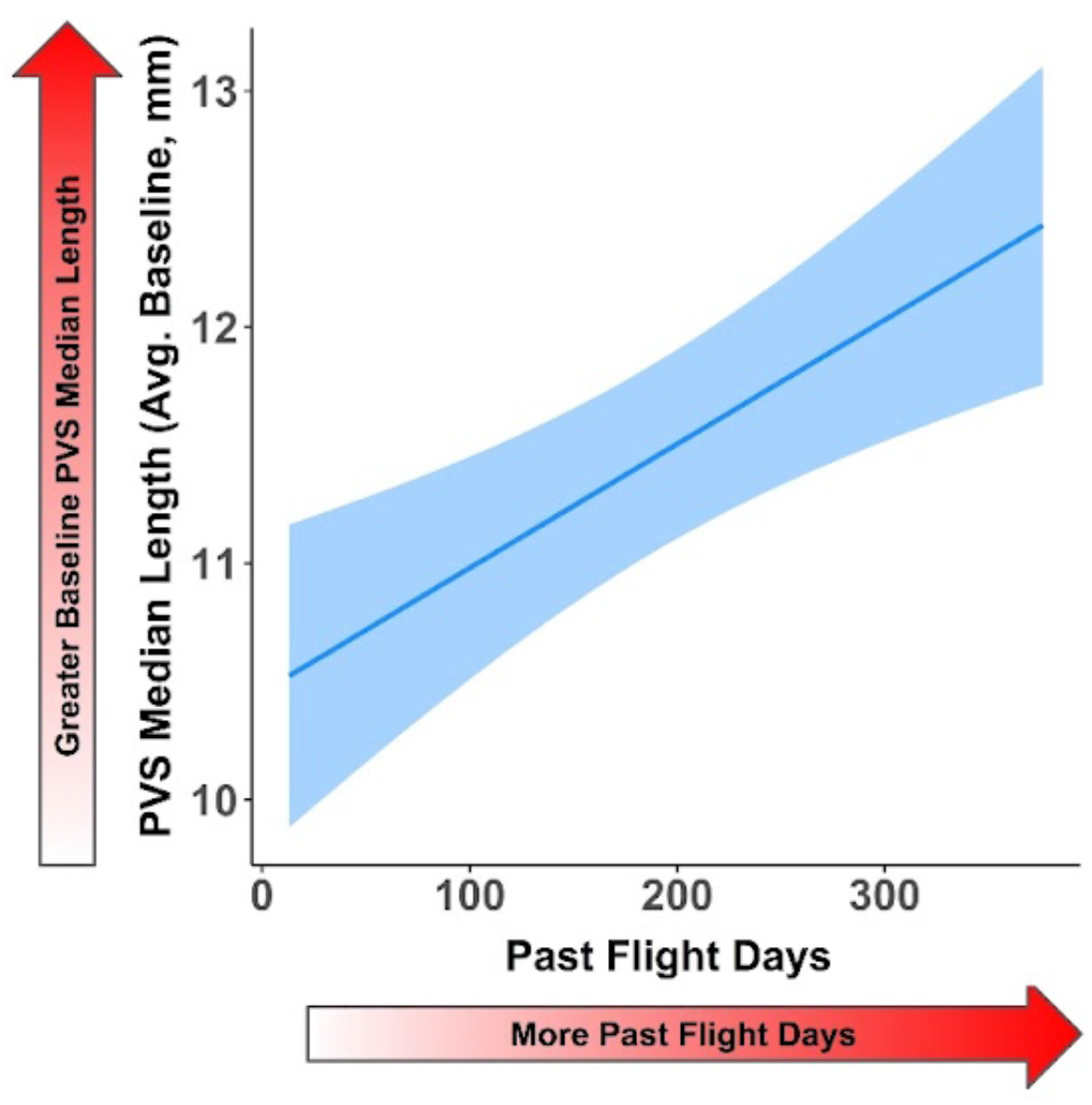
More Previous Days in Space Associates with Greater Pre-Flight PVS Length. This plot includes only the *n* = 6 experienced astronauts. Average pre-flight baseline (i.e., the mean of the two pre-flight time points) PVS median length (mm) is shown on the y-axis. Total number of past flight days is shown on the x-axis. To protect astronaut privacy, we do not show individual data points.

### 3.5 Correlation Between Post-Flight Ventricular Expansion and PVS Characteristics

The correlations between post-flight ventricular expansion and PVS characteristics are summarized in Table 4 and Fig. 4 (Table 4 shows all model effects, whereas unadjusted values are plotted in Fig. 4 for ease of interpretation). Pre- to post-flight change in right lateral ventricular volume was correlated with pre- to post-flight change in median PVS length. The stepwise variable selection procedure retained both right lateral ventricular volume (*p* = 0.008) and age at launch (*p* = 0.022) as significant predictors of pre- to post-flight change in PVS median length. Controlling for age, for every 1 percent increase in right lateral ventricular volume from pre- to post-flight, the median PVS length increased by 0.89 percent (*p* = 0.008). For completeness, we depict the identified relationship between older age at launch and greater pre- to post-flight changes in median PVS length in Fig. S1. There was no correlation between right lateral ventricular volume changes and the other PVS characteristics.

**Table 4.**
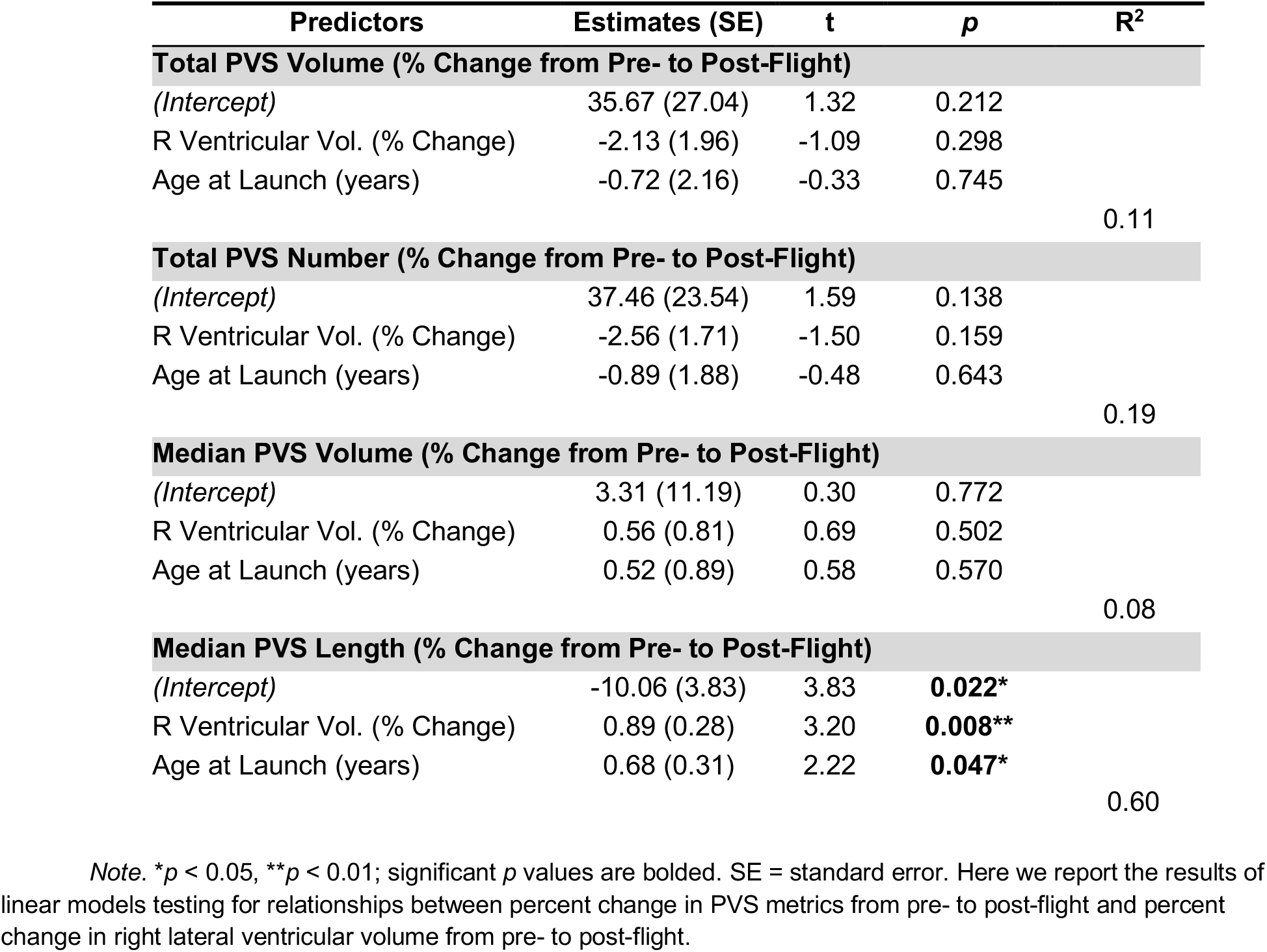
Correlation Between Post-Flight Ventricular Expansion and PVS Characteristics

**Fig 4.**
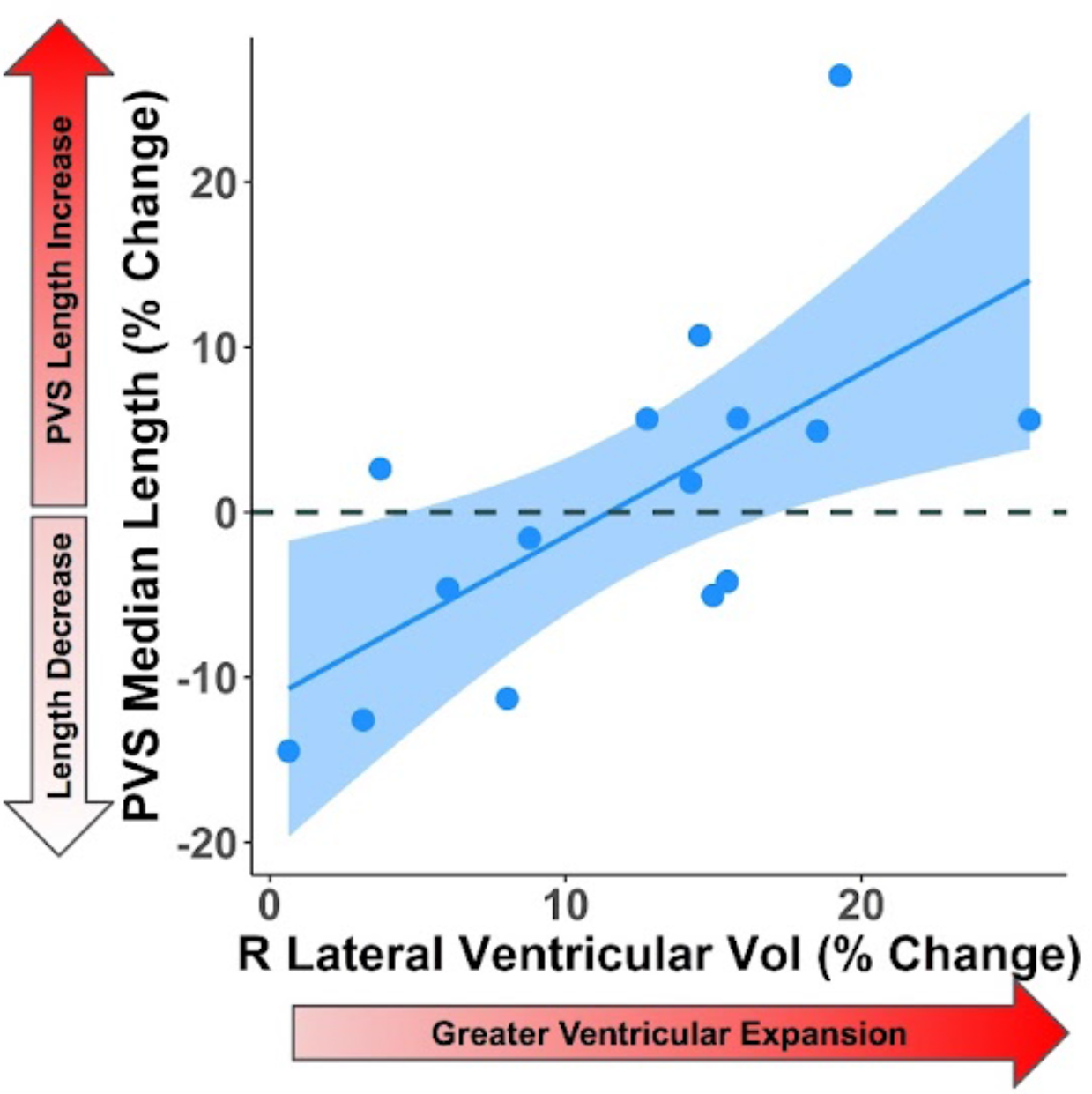
Association Between Greater Right Lateral Ventricular Volume Expansion and Pre- to Post-Flight Increases in PVS Median Length. Each point represents one astronaut. Right lateral ventricular volume % change from pre- to post-flight is plotted on the x-axis, and PVS median length % change from pre- to post-flight is plotted on the y-axis.

### 3.6 No Group Differences by SANS Status

There were no statistically significant changes in PVS metrics or right lateral ventricular volume at baseline from pre- to post-flight between those who did and did not experience one or more symptoms of SANS (Table S2).

## 4. Discussion

This is the first study to examine changes in MRI-visible PVS in the white matter with spaceflight. Our key findings included that novice astronauts exhibited increases in total PVS volume from pre- to post-flight, whereas experienced astronauts did not. Moreover, in experienced astronauts, the number of days spent in space prior to the current mission was associated with increased PVS length at our two pre-flight scan time points. This implies that the lack of PVS change with the current flight for this group was limited due to the effects of prior spaceflight missions on PVS. We also identified a significant, positive correlation between pre- to post-flight increases in PVS median length and increases in right lateral ventricular volume. Finally, we did not find any significant differences in PVS characteristics between those who did and did not exhibit SANS.

We found that novice astronauts exhibited an increase in PVS volume from pre- to post-flight, whereas experienced flyers did not. This divergence between novice and experienced astronauts is consistent with a compensatory hypothesis derived by Roberts and colleagues^45^, suggesting that spaceflight-induced ventricular changes are a compensatory response for cerebrospinal fluid hydrodynamics. That is, they suggest that experienced flyers have larger pre-flight ventricular volume (i.e., lingering ventricular expansion from previous flights^5,6,10,11^), and therefore do not show as much increase with subsequent spaceflights, perhaps due to a loss of compliance with changes from previous flights. Whether the morphological PVS findings observed here reflect pathophysiological or compensatory consequences of spaceflight remains unknown. Regardless, repeated spaceflight exposure appears to be a key determinant of brain fluid changes that should be further explored in the future. That is, these data suggest a cumulative and asymptotic effect on CSF spaces. As astronauts experience microgravity for longer cumulative durations, ventricles enlarge, and PVS size increases, eventually reaching a plateau, hence decreasing the changes observed with every subsequent flight.

In line with this hypothesis, we also found that, among the experienced crewmembers, PVS median length at baseline was correlated with the number of days previously spent in space. These outcomes follow previous work from our group and others, showing that alterations in gray matter^4,5^, white matter microstructure^8^ and hypointensities^12^, free water distribution^8^, and ventricular volume^3,5,9,12^ are dependent on the frequency and / or duration of time spent in space. This finding also suggests that PVS changes with spaceflight may be long lasting. Indeed, previous spaceflight longitudinal analyses have suggested that several structural brain changes with flight (namely, ventricular expansion) are slow to recover, remaining elevated at 6 to 12-months post-flight for some individuals ^5,6,10,11^. Therefore, as NASA’s goals shift towards longer-duration missions to Mars, it is becoming increasingly critical that we study the mechanisms and consequences of PVS enlargement in future spaceflight or ground-based analog studies, ideally with larger samples.

Furthermore, we observed a positive correlation between the percent increase in right lateral ventricular volume with flight and increases in PVS length. Previous studies have shown spaceflight-induced cranial fluid shifts^3,6,8^. These shifts likely compress the adjacent venous structures at the top of the head and obstruct the arachnoid granulations positioned along the superior sagittal sinus. It is also plausible that dural lymphatic vessels are compressed, further impairing CSF reabsorption. Roberts et al.^45^ have suggested that the result of this CSF outflow obstruction is ventricular dilation similar to that observed in individuals with normal pressure hydrocephalus. Under this assumption, the change in PVS morphology we observed in association with ventricular dilation may represent the result of CSF outflow obstruction. It is worth noting that individuals with normal pressure hydrocephalus show both dilated PVS and glymphatic dysfunction represented by delayed clearance of intrathecally administered gadolinium contrast^46–48^.

In our cohort, we did not observe an association between changes in PVS characteristics and clinical manifestations of SANS. Headward fluid shifts leading to obstruction of CSF and intraorbital fluid and glymphatic dysfunction have been proposed as potential mechanisms underlying SANS^13^. There are several plausible explanations for the lack of relationship between PVS change and SANS symptoms. First, it may be that PVS function does not relate to SANS. Second, the role of PVS in CSF clearance remains unknown, and it is possible that impaired clearance still occurs in the absence of PVS enlargement^19^. Third, the absence of correlations between PVS and SANS symptoms could be a consequence of our small subgroup sizes.

Our study has several limitations. The number of people who travel to space is small, creating an inevitable obstacle in the efforts to study spaceflight-associated changes. To address the concern of sample size, a repeated measures design was implemented, thereby allowing each astronaut to serve as their own control. There was also a delay between re-entry to Earth’s gravitational field and the first post-flight MRI acquisition, ranging between one and six days after landing (average = 4.5 days). We used this value as a covariate in analyses in an effort to mitigate its effect. Given that ventricular volume changes are still evident months post-flight^5,6,10,11^, this time offset is likely not a key factor in the current study. However, it could be that PVS and ventricular volume changes have different temporal dynamics and recovery patterns. Another limitation is that the PVS algorithm we employed is tailored to white matter PVS segmentation and has not yet been optimized for identifying PVS in the gray matter; future studies should enhance the algorithm to incorporate other tissue types and anatomical regions. It is possible that the PVS segmentation algorithm is not sensitive enough to uncover subtle differences in PVS characteristics caused by spaceflight. However, despite large between-subject differences, ICCs between the two pre-flight MRIs met our *a priori* inclusion criteria for all but one (i.e., median width) indicator of PVS characteristics. This finding argues in favor of our automated PVS detection algorithm’s reliability, as described in previous work. Due to the resolution of the images available, we were not able to distinguish whether the quantified PVSs were arterial or venular. Lastly, while our algorithm is able to detect PVS in *T*_1_-weighted images, the absence of a *T*_2_-weighted scan prevents PVS from being compared or contrasted with white matter hyperintensities, a condition potentially relevant to spaceflight^12^. Future studies using higher resolution images and multiple MR sequence types are likely to yield more information.

## 5. Conclusion

We found that at the group level, PVS was not affected by spaceflight. However, when we evaluated these changes separately between novice astronauts and those that had been to space before, we observed experience-dependent effects. First, novice astronauts had an increase in PVS with spaceflight, whereas experienced crewmembers did not. Second, those with prior spaceflight experience exhibited a correlation between more previous days in space and greater PVS length. Our findings suggest that PVS associations with spaceflight are dependent upon prior experience, similar to other brain structural changes we and others have reported^3–5,9,12,32^. Whether these changes in PVS represent a compensatory mechanism, or will eventually lead to pathological manifestations remains unknown. Given the relationship we identified between the percent increase in ventricular volume with flight and increases in PVS length, we suggest that our findings relate to impaired CSF outflow caused by the fluid^3,6,8^ and upward brain position shifts^3–5^ observed with microgravity exposure. However, we did not find an association between changes in PVS characteristics and clinical manifestations of SANS. Moving forward, head-down tilt bed rest analog studies may provide further insights into PVS changes, as this environment also results in brain position and fluid shifts.

## Acknowledgements

This work was supported by grants from the National Aeronautics and Space Administration (NASA NNX11AR02G) and the National Space Biomedical Research Institute (#NCC 9-58) to RS, AM, SW, and JB. During completion of this work, KH was supported by a National Science Foundation Graduate Research Fellowship under Grant no. DGE-1315138 and DGE-1842473, National Institute of Neurological Disorders and Stroke training grant T32-NS082128, and National Institute on Aging fellowship 1F99AG068440. DS was supported by the National Institute on Aging under Grant nos. R01AG056712 and P30AG008017. LS was supported by the National Institute on Aging under Grant no. P30AG066518. JP was supported by the National Heart Lung and Blood Institute under Grant no. K23HL150217-01. The authors also wish to thank all of the astronauts who volunteered their time, without whom this project would not have been possible.

## Author Contributions

KH extracted ventricular volumes, conducted all statistical analyses, created all figures and supplemental material, and wrote the Methods and Results. SBR, KH, RS, and JP wrote the Abstract, Introduction, and Discussion sections. HM conducted the SANS versus no-SANS classifications and participated in manuscript preparation. DS designed the PVS analysis pipeline and processed the images to extract PVS metrics. ML visually checked the results of the automated PVS pipeline. NB and YD collected and analyzed data. IS participated in project design and software development. RR facilitated data collection. SW, JB, AM, LS, JI, RS, and JP designed the project and led interpretation and discussion of the results. All authors participated in revision of the manuscript.

## References

1. Clément, G. R. et al. Challenges to the central nervous system during human spaceflight missions to Mars. Journal of Neurophysiology vol. 123 (2020).

2. Hupfeld, K. E., McGregor, H. R., Reuter-Lorenz, P. A. & Seidler, R. D. Microgravity effects on the human brain and behavior: Dysfunction and adaptive plasticity. Neuroscience and Biobehavioral Reviews vol. 122 (2021).

3. Roberts, D. R. et al. Effects of Spaceflight on Astronaut Brain Structure as Indicated on MRI. New England Journal of Medicine 377, (2017).

4. Koppelmans, V., Bloomberg, J. J., Mulavara, A. P. & Seidler, R. D. Brain structural plasticity with spaceflight. npj Microgravity 2, (2016).

5. Hupfeld, K. E. et al. The impact of six and twelve months in space on human brain structure and intracranial fluid shifts. Cerebral Cortex Communications 1, (2020).

6. Jillings, S. et al. Macro-And microstructural changes in cosmonauts’ brains after long-duration spaceflight. Science Advances 6, (2020).

7. Riascos, R. F. et al. Longitudinal Analysis of Quantitative Brain MRI in Astronauts Following Microgravity Exposure. Journal of Neuroimaging 29, (2019).

8. Lee, J. K. et al. Spaceflight-Associated Brain White Matter Microstructural Changes and Intracranial Fluid Redistribution. in JAMA Neurology vol. 76 (2019).

9. Roberts, D. R. et al. Prolonged microgravity affects human brain structure and function. American Journal of Neuroradiology 40, (2019).

10. Kramer, L. A. et al. Intracranial effects of microgravity: A prospective longitudinal MRI study. Radiology 295, (2020).

11. van Ombergen, A. et al. Brain ventricular volume changes induced by long-duration spaceflight. Proceedings of the National Academy of Sciences of the United States of America 116, (2019).

12. Alperin, N., Bagci, A. M. & Lee, S. H. Spaceflight-induced changes in white matter hyperintensity burden in astronauts. Neurology 89, (2017).

13. Lee, A. G. et al. Spaceflight associated neuro-ocular syndrome (SANS) and the neuro-ophthalmologic effects of microgravity: a review and an update. npj Microgravity vol. 6 (2020).

14. Lee, A. G., Mader, T. H., Gibson, C. R. & Tarver, W. Space flight-associated neuro-ocular syndrome. JAMA Ophthalmology 135, (2017).

15. Iliff, J. J. et al. Brain-wide pathway for waste clearance captured by contrast-enhanced MRI. Journal of Clinical Investigation 123, (2013).

16. Iliff, J. J. et al. A paravascular pathway facilitates CSF flow through the brain parenchyma and the clearance of interstitial solutes, including amyloid β. Science Translational Medicine 4, (2012).

17. Louveau, A. et al. Structural and functional features of central nervous system lymphatic vessels. Nature 523, (2015).

18. Bacyinski, A., Xu, M., Wang, W. & Hu, J. The paravascular pathway for brain waste clearance: Current understanding, significance and controversy. Frontiers in Neuroanatomy vol. 11 (2017).

19. Brown, R. et al. Understanding the role of the perivascular space in cerebral small vessel disease. Cardiovascular Research vol. 114 (2018).

20. Ramirez, J. et al. Visible Virchow-Robin spaces on magnetic resonance imaging of Alzheimer’s disease patients and normal elderly from the Sunnybrook dementia study. Journal of Alzheimer’s Disease 43, (2015).

21. Potter, G. M., Chappell, F. M., Morris, Z. & Wardlaw, J. M. Cerebral perivascular spaces visible on magnetic resonance imaging: Development of a qualitative rating scale and its observer reliability. Cerebrovascular Diseases 39, (2015).

22. Charidimou, A. et al. White matter perivascular spaces: An MRI marker in pathology-proven cerebral amyloid angiopathy? Neurology 82, (2014).

23. Inglese, M. et al. Clinical significance of dilated Virchow-Robin spaces in mild traumatic brain injury. Brain Injury 20, (2006).

24. Patankar, T. F. et al. Dilatation of the Virchow-Robin space is a sensitive indicator of cerebral microvascular disease: Study in elderly patients with dementia. American Journal of Neuroradiology 26, (2005).

25. Opel, R. A. et al. Effects of traumatic brain injury on sleep and enlarged perivascular spaces. Journal of Cerebral Blood Flow and Metabolism 39, (2019).

26. Hupfeld, K. E. et al. Brain and Behavioral Evidence for Reweighting of Vestibular Inputs with Long-Duration Spaceflight. Cerebral Cortex (2021) doi:10.1093/cercor/bhab239.

27. Tays, G. et al. The effects of long duration spaceflight on sensorimotor control and cognition. bioRxiv (2021).

28. Koppelmans, V. et al. Study protocol to examine the effects of spaceflight and a spaceflight analog on neurocognitive performance: Extent, longevity, and neural bases. BMC Neurology 13, (2013).

29. Dahnke, R., Yotter, R. A. & Gaser, C. Cortical thickness and central surface estimation. NeuroImage 65, (2013).

30. Gaser, C. & Kurth, F. Manual computational anatomy toolbox-CAT12. (Structural brain mapping Group at the Departments of Psychiatry and Neurology, University of Jena, 2017).

31. Ashburner, J. et al. SPM12 manual. vol. 2464 (Wellcome Trust Centre for Neuroimaging, 2014).

32. Lee, J. K. et al. Effects of Spaceflight Stressors on Brain Volume, Microstructure, and Intracranial Fluid Distribution. Cerebral Cortex Communications 2, (2021).

33. Scotti, G. et al. MR imaging of cavernous sinus involvement by pituitary adenomas. American Journal of Neuroradiology 9, (1988).

34. Ayanzen, R. H. et al. Cerebral MR venography: Normal anatomy and potential diagnostic pitfalls. American Journal of Neuroradiology 21, (2000).

35. Mader, T. H. et al. Persistent Asymmetric Optic Disc Swelling after Long-Duration Space Flight: Implications for Pathogenesis. Journal of Neuro-Ophthalmology 37, (2017).

36. Piantino, J. et al. Link between Mild Traumatic Brain Injury, Poor Sleep, and Magnetic Resonance Imaging: Visible Perivascular Spaces in Veterans. Journal of Neurotrauma (2021) doi:10.1089/neu.2020.7447.

37. Piantino, J. et al. Characterization of MR imaging-visible perivascular spaces in the white matter of healthy adolescents at 3T. American Journal of Neuroradiology 41, (2020).

38. Schwartz, D. L. et al. Autoidentification of perivascular spaces in white matter using clinical field strength T1 and FLAIR MR imaging. NeuroImage 202, (2019).

39. Promjunyakul, N. et al. Characterizing the white matter hyperintensity penumbra with cerebral blood flow measures. NeuroImage: Clinical 8, (2015).

40. Team, T. R. D. C. The R Reference Manual. Development 1, (2004).

41. Revelle, W. Package “psych” - Procedures for Psychological, Psychometric and Personality Research. R Package (2015).

42. Koppelmans, V. et al. Brain plasticity and sensorimotor deterioration as a function of 70 days head down tilt bed rest. PLoS ONE 12, (2017).

43. Koo, T. K. & Li, M. Y. A Guideline of Selecting and Reporting Intraclass Correlation Coefficients for Reliability Research. Journal of Chiropractic Medicine 15, (2016).

44. Venables, W. N. & Ripley, B. D. Modern Applied Statistics with S-Plus. Biometrics 52, (1996).

45. Roberts, D. R. & Petersen, L. G. Studies of Hydrocephalus Associated with Long-term Spaceflight May Provide New Insights into Cerebrospinal Fluid Flow Dynamics Here on Earth. JAMA Neurology vol. 76 (2019).

46. Ringstad, G., Vatnehol, S. A. S. & Eide, P. K. Glymphatic MRI in idiopathic normal pressure hydrocephalus. Brain 140, (2017).

47. Ringstad, G. et al. Brain-wide glymphatic enhancement and clearance in humans assessed with MRI. JCI insight 3, (2018).

48. Eide, P. K., Pripp, A. H., Ringstad, G. & Valnes, L. M. Impaired glymphatic function in idiopathic intracranial hypertension. Brain Communications 3, (2021).

