## Supplementary material for "MRI-Visible Perivascular Space (PVS) Changes with Long-Duration Spaceflight": Table S1

**Table S1**. PVS Changes and Ventricular Expansion from Pre- to Post-Flight

| Predictors | Estimates (SE) | t | *p* | $R^2$ |
|---|---|---|---|---|
| **Total PVS Volume (% Change)** | | | | |
| *(Intercept)* | 11.16 (16.73) | 0.69 | 0.505 | |
| Age | -1.29 (2.47) | -0.52 | 0.613 | |
| Sex | -5.41 (34.07) | -0.16 | 0.877 | |
| Landing to MRI Time | 2.50 (11.17) | 0.22 | 0.827 | |
| | | | | 0.03 |
| **Total PVS Number (% Change)** | | | | |
| *(Intercept)* | 6.03 (15.08) | 0.40 | 0.697 | |
| Age | -1.42 (2.23) | -0.64 | 0.536 | |
| Sex | 0.87 (30.71) | 0.03 | 0.978 | |
| Landing to MRI Time | 4.24 (10.07) | 0.42 | 0.682 | |
| | | | | 0.05 |
| **Median PVS Volume (% Change)** | | | | |
| *(Intercept)* | 10.58 (6.76) | 1.57 | 0.146 | |
| Age | 0.58 (1.00) | 0.58 | 0.574 | |
| Sex | -1.74 (13.76) | -0.13 | 0.902 | |
| Landing to MRI Time | -0.02 (4.51) | -0.01 | 0.996 | |
| | | | | 0.04 |
| **Median PVS Length (% Change)** | | | | |
| *(Intercept)* | 2.48 (2.91) | 0.85 | 0.413 | |
| Age | 0.70 (0.43) | 1.61 | 0.135 | |
| Sex | -6.59 (5.93) | -1.11 | 0.291 | |
| Landing to MRI Time | -1.00 (1.95) | -0.52 | 0.617 | |
| | | | | 0.34 |
| **Right Lateral Ventricular Volume (% Change)** | | | | |
| *(Intercept)* | 12.60 (2.29) | 5.49 | **0.0002**** | |
| Age | 0.16 (0.34) | 0.48 | 0.642 | |
| Sex | -1.69 (4.67) | -0.36 | 0.724 | |
| Landing to MRI Time | -1.16 (1.53) | -0.76 | 0.465 | |

*Table S1 Note*. **$p < 0.01$; significant *p* values are bolded. SE = standard error. Here we report the results of linear models testing whether the percent change in each PVS metric and right lateral ventricular volume from pre- to post-flight differed significantly from 0, controlling for age at launch, sex, and the time between landing and the first post-flight time point. Males served as the reference group (i.e., coded as = 0) for the categorical sex variable. Our primary interest here was whether the intercept was significant ($p < 0.05$), thereby indicating a significant whole-group change in the PVS metric or ventricular volume with spaceflight.

**Fig. S1**. Association Between Older Age at Launch and Greater Pre- to Post-Flight Increases in PVS Median Length

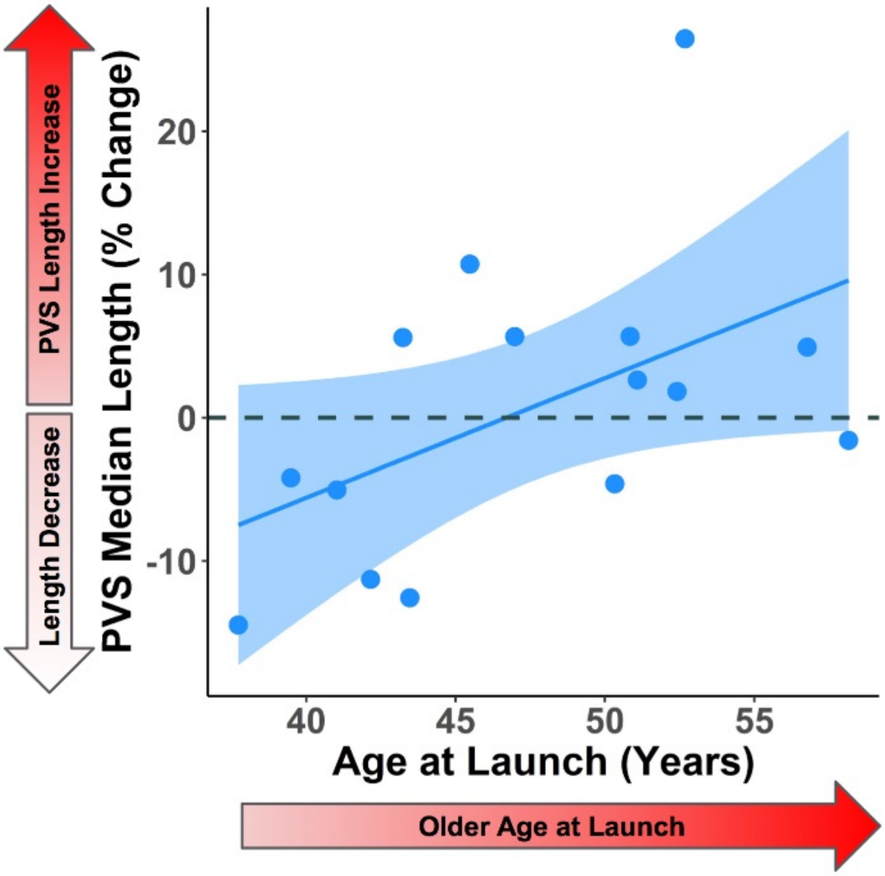


*Fig. S1 Note*. Each point represents one astronaut. PVS median length % change from pre- to post-flight is plotted on the y-axis, and age at launch (years) is plotted on the x-axis. We provide this plot for completeness, as age at launch was one of the two retained significant predictors of PVS median length changes with flight. (The other significant predictor of PVS median length changes with flight was right lateral ventricular volume expansion; see Fig. 4 and Table 4 in the main text.)

**Table S2**. PVS Changes and Ventricular Expansion from Pre- to Post-Flight: SANS vs. no-SANS

| Predictors | Estimates (SE) | t | *p* | $R^2$ |
|---|---|---|---|---|
| **Total PVS Volume (% Change)** | | | | |
| *(Intercept)* | 30.02 (29.91) | 1.00 | 0.349 | |
| SANS vs. no-SANS | -27.62 (41.35) | -0.67 | 0.526 | |
| Age | -0.23 (3.14) | -0.07 | 0.944 | |
| Sex | -13.51 (30.87) | -0.44 | 0.675 | |
| Landing to MRI Time | 2.98 (10.16) | 0.29 | 0.778 | |
| | | | | 0.13 |
| **Total PVS Number (% Change)** | | | | |
| *(Intercept)* | 25.54 (30.95) | 0.83 | 0.437 | |
| SANS vs. no-SANS | -23.17 (42.79) | -0.54 | 0.605 | |
| Age | -0.30 (3.25) | -0.09 | 0.929 | |
| Sex | -8.76 (31.95) | -0.27 | 0.792 | |
| Landing to MRI Time | 4.78 (10.52) | 0.46 | 0.663 | |
| | | | | 0.11 |
| **Median PVS Volume (% Change)** | | | | |
| *(Intercept)* | 4.70 (9.57) | 0.49 | 0.639 | |
| SANS vs. no-SANS | -3.35 (13.23) | -0.25 | 0.807 | |
| Age | -0.05 (1.01) | -0.05 | 0.965 | |
| Sex | 3.53 (9.88) | 0.36 | 0.731 | |
| Landing to MRI Time | 2.61 (3.25) | 0.80 | 0.449 | |
| | | | | 0.12 |
| **Median PVS Length (% Change)** | | | | |
| *(Intercept)* | 2.04 (5.26) | 0.39 | 0.710 | |
| SANS vs. no-SANS | -3.53 (7.28) | -0.49 | 0.642 | |
| Age | 0.71 (0.55) | 1.28 | 0.240 | |
| Sex | -5.00 (5.43) | -0.92 | 0.388 | |
| Landing to MRI Time | -0.27 (1.79) | -0.15 | 0.886 | |
| | | | | 0.34 |
| **Right Lateral Ventricular Volume (% Change)** | | | | |
| *(Intercept)* | 12.60 (5.76) | 2.19 | 0.065 | |
| SANS vs. no-SANS | -1.57 (7.96) | -0.20 | 0.849 | |
| Age | 0.18 (0.61) | 0.30 | 0.770 | |
| Sex | -1.17 (5.94) | -0.20 | 0.850 | |
| Landing to MRI Time | -0.98 (1.96) | -0.50 | 0.631 | |
| | | | | 0.05 |

*Table S2 Note.* SE = standard error. Here we report the results of linear models testing whether the percent change in each PVS metric and right lateral ventricular volume from pre- to post-flight differed for the SANS vs. no-SANS astronauts, controlling for age at launch, sex, and the time between landing and the first post-flight MRI scan. no-SANS and male served as the reference groups (i.e., coded as = 0) for the two categorical variables. Our primary interest here was whether there was an effect of SANS status on pre- to post-flight percent change in the PVS and ventricle metrics.
